# Millisecond-scale behaviours of plankton quantified *in situ* and *in vitro* using the Event-based Vision Sensor (EVS)

**DOI:** 10.1101/2023.01.11.523686

**Authors:** Susumu Takatsuka, Norio Miyamoto, Hidehito Sato, Yoshiaki Morino, Yoshihisa Kurita, Akinori Yabuki, Chong Chen, Shinsuke Kawagucci

## Abstract

The Event-based Vision Sensor (EVS) is a bio-inspired sensor that captures detailed motions of objects, developed with the applicability to become the ‘eyes’ of machines and especially self-driving cars. Compared to conventional frame-based image sensors as employed in video cameras, EVS has an extremely fast motion capture equivalent to 10,000-fps even with standard optical settings and additionally has high dynamic ranges for brightness and also lower consumption of memory and energy. These features make the EVS an ideal method to tackle questions in biology, such as the fine-scale behavioural ecology. Here, we developed 22 characteristic features for analysing the motions of aquatic particles from the raw data of the EVS, and deployed the EVS system in both natural environments and laboratory aquariums to test its applicability to filming and analysing plankton behaviour. Our EVS monitoring in turbid water at the bottom of Lake Biwa, Japan identified several particles exhibiting distinct cumulative trajectory with periodicities in their motion (up to 16 Hz), suggesting that they were living organisms with rhythmic behaviour. We also carried out EVS monitoring in the deep sea aided by infrared lighting to minimise influence on behaviour, and observed particles with active motion and periodicities over 40 Hz. Furthermore, we used the EVS to observe laboratory cultures of six species of zooplankton and phytoplankton, confirming that they have species-specific motion periodicities of up to 41 Hz. We applied machine learning to automatically classify particles into five categories (four categories of zooplankton plus passive particles), which achieved an accuracy up to 86%. Our attempts to use the EVS for biological observations, especially focusing on its millisecond-scale temporal resolution and wide dynamic range provide a new avenue to investigate rapid and periodical motion and behaviour in small organisms. Given its compact size with low consumption of battery and memory, the EVS will likely be applicable in the near future for the automated monitoring of the behaviour of plankton by edge computing on autonomous floats, as well as quantifying rapid cellular-level activities under microscopy.

## 1. Introduction

Visual observation is a fundamental piece of information for biology. Human beings have been documenting the visual information of species, such as their general appearance and behaviour, from the early history evident in cave art and murals (Brumm et al., 2021). Recording of organismal behaviour and morphology went from drawings to photography and, in the last century, video cameras were developed. Videos allow people to objectively capture organisms’ behaviour, and opened avenues of research into the functions and motions of individual bodies and body parts, spatiotemporal tracking of individuals in the natural environment, and physical-social interactions (e.g. grazing, mating, etc.) among individuals and species. In addition to qualitative descriptions based on video recordings, quantitative data extracted by mathematical analysis enabled us to objectively interpret and understand the behaviour of species. Typically, videos are frame-based, meaning they are a series of still images where information from all pixels (i.e. individual sensors corresponding to a pixel) on the image sensor is recorded synchronously as a ‘frame’.

High-speed video imaging is a technology allowing the recording of much more frames per unit time than what the human eye can normally perceive. While human vision image acquisition is no faster than the order of 100 frames per second (fps) (Potter et al., 2014), high-speed videos have achieved >1,000 fps (one frame per millisecond) (e.g., Thoroddsen et al., 2008) revolutionising visual observations of biological phenomena. For example, high-speed video analyses revealed quantitative data and trends in the behaviour of plankton, such as changes in speed/acceleration/orientation at the millisecond-scale (Buskey et al., 2002). However, the current high-speed video imaging techniques are still frame-based, and since shorter exposure time means less light per frame per photosite, there is a demand for strong lighting to achieve millisecond timeframe. This also means the system consumes a high amount of power, and can only be done over a short duration due to heavy memory consumption (Table 1). This strong lighting required for frame-based highspeed imaging can also influence animal behaviour and impact the results. The size of these commercially available high-speed video systems makes them non-portable and are cumbersome and costly to apply to natural ecosystem monitoring.

**Table 1.** A comparison table with Gopro Hero 8, FASTCAM Nova S6 and EVS prototype.

|  | Gopro Hero 8 | FASTCAM Nova S6 | EVS prototype |
| --- | --- | --- | --- |
| Resolution | 1280×820 | 1024x768 | 1280x720 |
| Flame rate | 240fps | 9000fps | 10000fps<br>Equivalent temporal p<br>recision |
| Data size per minute | 340 MB<sub>(MP4)</sub> | 594 GB | 200~500 MB |
| Power consumption with memory and peripheral circuits | 1.2~1.5W | 138W*<sub>1</sub> | 10.2~12.5W*<sub>2</sub> |
| Light power consumption | 33W*<sub>3</sub> | 120W*<sub>4</sub> | 1W |

\*1 https://www.webj.co.jp/wp-content/uploads/2019/01/7aec9ebad50b2718d72a0c3b411b6234.pdf
\*2 EVS prototype camera including FPGA and M.2 memory
\*3 Edoko light power consumption
\*4 https://www.h-repic.co.jp/products/fiber/fiber_led/la-hdf_8010

The Event-based Vision Sensor, EVS (Gallego et al., 2022; Lichtsteiner et al., 2008), also known as event camera, dynamic vision sensor, neuromorphic camera, or silicon retina, is a bio-inspired sensor, differs from frame-based systems in that the information in each pixel is recorded separately, when there is a change in brightness (Fig. 1A–C). Each pixel records a different output depending on the polarity of the brightness change. For example, an increase in brightness is recorded as a positive (+) change, while a decrease in brightness is recorded as a negative (-) change (Fig. 1D). These changes are called *events*, hence the name EVS. The EVS generates a dataset consisting of coordinates (X/Y), polarities (+/-), and timestamps of *events* in each pixel – meaning there is no need to record information from those pixels with no changes over time. This allows a much reduced data size as well as memory and power consumption, meaning it can track the movement of objects much more efficiently under lower light conditions while having less power and memory consumption (Table 1). The resulting four-dimensional dataset can be directly used to analyse the motion of objects, but the same data can also be used to simulate the recorded changes visually for cross-checking the object’s tracks with human eyes (Fig. 1D). The file size is two or more magnitudes smaller, while the power consumption is an order of magnitude less than frame-based high-speed video cameras (e.g. FASTcam Nova S6) currently on the market. Since the commercial distribution of the EVS began in 2008 (Lichtsteiner et al., 2008), it has been anticipated as a solution to improved computer vision with applicability to devices such as a self-driving car (Maqueda et al., 2018). These attributes of the EVS, however, mean that it also has great applicability to observe and analyse the detailed movement and behavioural patterns of organisms.

**Figure 1.**
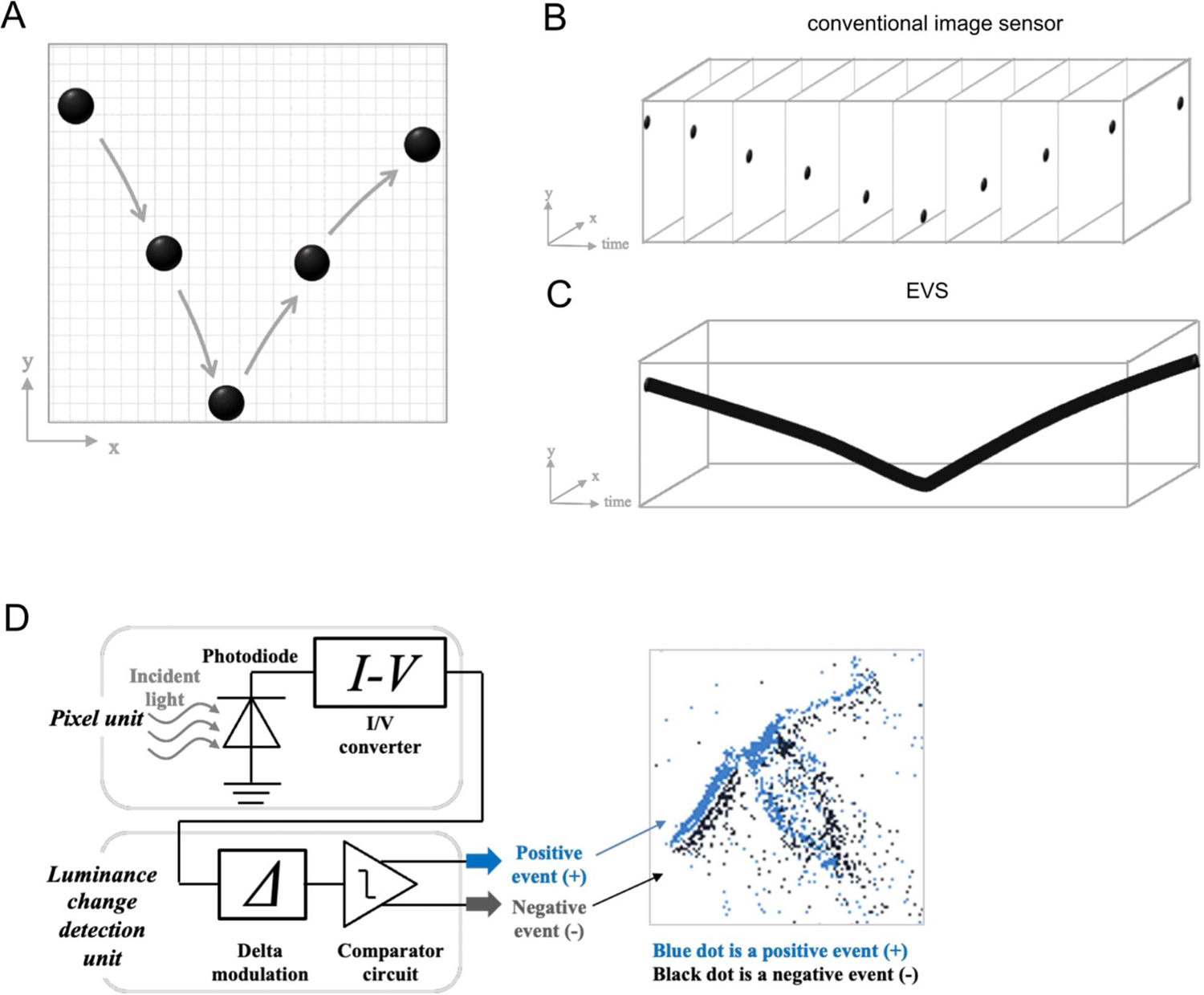
General Introduction of EVS. **A–C.** Differences between a conventional frame-based image sensor and the EVS when recording a bounding ball. **B.** Frame-based image sensors record all pixel information together, in ‘frames’. **C.** The EVS detects brightness changes asynchronously at each pixel and only those pixels with changes are recorded. **D.** The mechanism of an EVS as a circuit diagram and output image.

Here, we report the first application of the EVS to observing and analysing the behaviour of plankton in the aquatic environment. Living plankton are representative ‘active particles’ in natural aquatic systems, and *in situ* behaviours of the living organisms and their ecological functions are central issues of aquatic biology as well as biological limnology and oceanography (e.g. McManus and Woodson, 2012). At the same time, numerous ‘inactive particles’ such as dead bodies, fecal pellets, minerals, and their aggregates are floated by matrix liquid flow and gravitated toward the floor. The EVS system protected by water-proof (and pressure-tolerant for deep water) housings successfully identified swimming behaviours of plankton even in dense particle storms and marine snow at the bottoms of both a lake and an ocean. Testing the system mounted onto a microscope in a controlled laboratory aquarium setting shows that the system can automatically distinguish three zooplankton species among mixed suspended particles. Our results demonstrate strong potential for the EVS to be applied to life science research, beyond industrial applications.

## 2. Materials and settings

### 2.1 Sensor and camera systems

We used a stacked Event-based Vision Sensor (IMX636, manufactured by Sony Semiconductor Solutions Corporation) with the resolution of 1280 × 720 pixels and 4.86 μm × 4.86 μm individual pixel size. In this study, the same model of EVS sensor was fitted on two different camera systems to suit *in vitro* and *in situ* observations. For *in vitro* observation, we used an EVS prototype camera system, a developer’s model that was controlled and powered via USB-C using a Windows laptop. For *in situ* observation in lacustrine and deep-sea environments, we modified the EVS prototype camera system to prepare a stand-alone EVS camera system that did not need to be connected to a laptop. The stand-alone EVS camera system consisted of the EVS (IMX636, Sony), a field-programmable gate array (FPGA) (XCZU3EG-2SFVC784, Xilinx), an Arduino board (Arduino Pro Mini 3.3V 8MHz, Aideepen), an M.2 memory card (SB-RKTQ-8TB, SABRENT), and Bluetooth peripheral circuits (sh-hc-06, DSD TECH). This *in situ* EVS observation system was operated by an external power supply through the USB Power Delively standard and a 25,600-mAh portable battery that allowed >6 hours of continuous operation. Optical interface of both camera systems was developed for the CS-mount lens standard.

### 2.2 Settings of the camera systems

In order to obtain complementary images between an EVS camera and a conventional frame-based camera for the evaluation of EVS performance, we developed a recording system. For *in situ* observations in natural aquatic environments, a cubic beam splitter (ELIOTEC CORP.) was placed behind an objective lens to provide the same images to both the EVS system and the conventional camera (GoPro Hero 8 or Hero 7, modified to C-mount, GoPro Inc., Supplementary Fig. S1). For *in vitro* laboratory work, both the EVS system and a conventional camera (HOZAN USB Camera L-835) were mounted on a stereomicroscope (YS05T, Micronet Inc.) and the optical axis from the same objective lens was split to take complementary videos in both systems.

For *in situ* observation at lake system, the EVS camera system with a Variable Focus Iris Lens (C-Mount, Focal length: 4-12mm, Aperture setting: F16-22) was contained in a glass sphere mounted on the stand-alone monitoring system *Edokko Mark 1* model COEDO (Kawagucci et al., 2020) (see Section 4.1, Supplementary Fig. S2). The COEDO provides white light for conventional frame-based video imaging. For *in situ* observation in the deep-sea environment, the EVS camera system with a Variable Focus Iris Lens (C-Mount + 1mm Spacer Ring, Focal length: 4-12mm, Aperture setting: F16-22) was contained in a 35 MPa-tolerant pressure housing made with aluminum alloy (Japan Industrial Standard, A7075) coated by TUFRAM^(R)^ with a 100 mm diameter viewport of 50 mm thick transparent acrylic cone (see Section 4.2, Supplementary Fig. S3). To constrain the depth of field and exclude distant objects that will be out of focus, a black screen was placed at 20 or 40 mm distance from the external surface of the acrylic window (Supplementary Fig. S3C, F). Eight infrared LED lights (850 nm wavelength) with the total electric power of 1 W (Model number: OPTOSUPPLY OSI3XNE3E1E) were configured in a circle around the objective lens in the pressure housing (Supplementary Fig. S3E) to provide illumination in the lightless deep sea, while minimising disruption to biological activities.

### 2.3 *In situ* evaluations

In order to evaluate the applicability of the EVS system to observing planktic organisms in natural aquatic systems, we attempted to record and analyse the behaviour of particles in Lake Biwa and Suruga Bay, Japan. For Lake Biwa, the EVS system mounted on *Edokko Mark 1* model COEDO (Supplementary Fig. S2) was deployed at the bottom of Lake Biwa at a depth of 60 m on March 8, 2022 (Supplementary Fig. S4A, B, Yamada et al., 2021). The main body of *Edokko Mark 1* COEDO was buoyant from the glass spheres, and so was anchored by a ballast and rope during the deployment. The altitude of the EVS camera when deployed was ca. 1 m from the lake floor. The water temperature *in situ* was approximately 6.8–8.5 °C. Recording was conducted during the daytime from 11:36 to 20:22 (8h 46m) Japan standard time (JST) under sunlight with additional lighting from the COEDO main body. The lake water appeared cloudy at the surface, due to turbid inflows from neighbouring rivers due to heavy snow falls several days prior to the observation.

Further to the lake testing, to test the applicability of the EVS’ wide dynamic range which enabled detection of subtle differences in the low luminance range, the EVS system was installed into a 35 MPa-tolerant housing and tested in total darkness in the deep sea (Supplementary Fig. S3A–D). The deep-sea EVS system was secured on the rear side of the Remotely Operated Vehicle (ROV) *Kaiko* (with vehicle *Mk-IV*) facing the water column, on a dive to 1,236 m depth in Suruga Bay, Japan during the last research cruise of the now-retired R/V *Kairei* (KR21-17) on January 24, 2022 (Supplementary Fig. S3F–H and S4A, C, Kawagucci et al., 2020). The altitude of the EVS from the seafloor was approximately 1 m, where the water temperature was 3 °C. Recording was conducted with infrared LEDs equipped in the pressure-tolerant housing (Supplementary Fig. S3E), in addition to the front-facing LEDs for the ROV’s main video cameras. The total recording time of the EVS was 3 hours, from 10:00 to 13:00 JST.

### 2.4 *In vitro* evaluation

For the *in vitro* evaluation of the EVS sensor, we used four species of zooplankton and two species of marine algae and observed them with an EVS system in the laboratory. The paracalanid marine copepod *Parvocalanus crassirostris* was purchased from Pacific Trading Co., Ltd. and reared in a 10 L tank. Eggs of the brine shrimp *Artemia* sp. were purchased from Kyorin Co., Ltd., and hatched by aeration in seawater. Sexually mature individuals of the true limpet *Nipponacmea fuscoviridis* were collected from Hiraiso, Ibaraki Prefecture and Tsuyazaki, Fukuoka Prefecture in Japan; mature individuals of the sea star *Patiria pectinifera* were collected from Tsuyazaki and Asamushi, Aomori Prefecture in Japan. These two species were reared in aquaria and *in vitro* fertilization was used to obtain their larvae, as previously described (Hashimoto et al., 2012; Koga et al., 2010). Two phytoplankton species, *Chattonella marina* (NIES-1) and *Heterosigma akashiwo* (NIES-5), were provided by National Institute for Environmental Studies (NIES). through the National BioResource Project (NBRP) of the Ministry of Education (MEXT), Japan.

During observation, the stereomicroscope was tilted 90 degrees because the target movements were the vertical movement of swimming plankton as well as particles settling under the influence of gravity (Supplementary Fig. S5A, B). A narrow acrylic aquarium with an inner thickeness of 5 mm between the walls was placed in front of the object lens for observation (Supplementary Fig. S5C). Several dozen individuals of the target organism were introduced into the aquarium, and the movement of *particles* was recorded. The behaviour of each plankton species was observed by EVS and the output analysed. Three zooplankton species, including *Parvocalanus crassirostris*, *N. fuscoviridis*, and *Patiria pectinifera*, were recorded together in a random mixture to test the capacity of the computer software to tease apart the species.

### 2.5 Development of data processing methods: particle identification

We developed a software [-to be published on Sony Group Corporation GitHub]to analyse the EVS outputs, consisting of pixel coordinates (X/Y), polarities in the change of brightness (+/-), and timestamps of *events*, asynchronously recorded from each independent pixel. Since the dataset generated by the EVS is quite different from conventional frame-based cameras, it is difficult to use conventional image processing software libraries such as OpenCV for the analyses of EVS data. Analyses of the EVS outputs can be conducted simply based on the recorded raw data without producing any visual images, but a pseudo-frame-based image can be simulated by accumulating the EVS outputs within a short-time interval for the purpose of visual confirmation by human eyes.

Densely aggregated, simultaneous local *events* within a short period of time that are isolated from other distant *events*, are recognized as a *cluster*, generally corresponding to a physical *particle* in the ecosystem observed. (Fig. 2A). Each *cluster* is composed of *events* which occurred less than 100 msec ago and all *events* older than that period were removed in order to track each *cluster’s* movement. For each new *event*, a region of 5×5 pixels around the new *event* was searched to decide which *cluster* the new *event* should be merged into (Fig. 2B). Iteration of this process leads to the growth of each *cluster*, until the shape of the *cluster* matches that of the particle (Fig. 2C). Temporal shifts of the absolute positions of a *cluster*, which means the combination between removing older *events* and adding new *events* in a specific timeframe, is recognized as the movement of a particle (Fig. 2D). An example of a reconstructed visual simulated by accumulating outputs across a 100 msec timeframe is shown in Fig 2E. Each *cluster* recognized is indicated by a rectangular boundary and assigned to a specific number.

**Figure 2.**
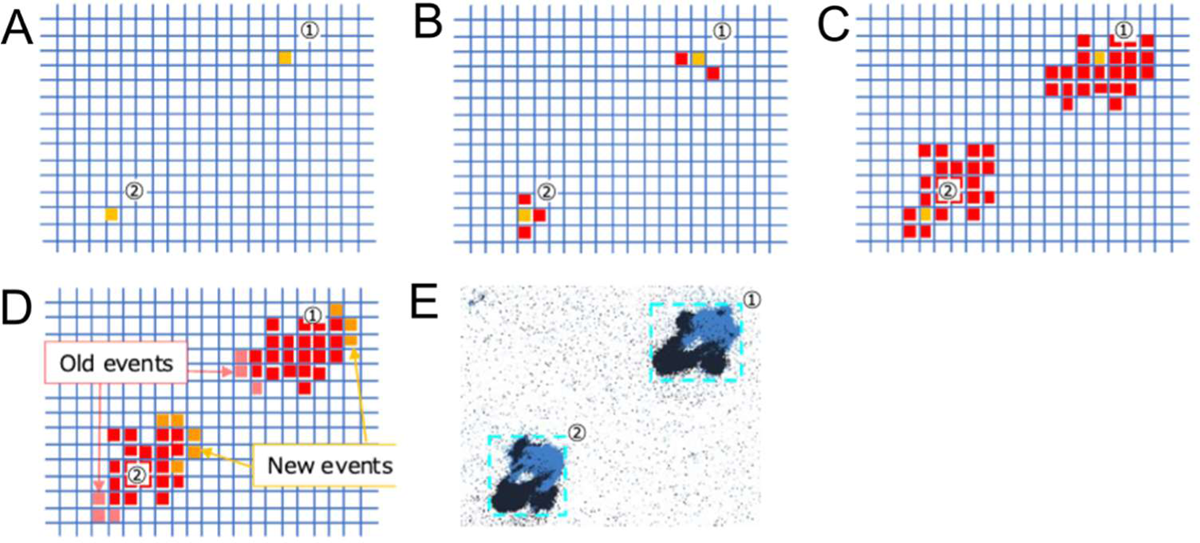
Recognition of particles and the conversion to *clusters*. **A.** Labelling of isolated *events* with *cluster* IDs. **B.** New *events* close to an existing *cluster* are assigned to the *cluster*. **C.** By repeating the process of **B**, the *cluster* eventually takes the form of a particle observed. **D.** By deleting old *events* and adding new *events*, the *cluster’s* motion is captured and allows for the tracking of the particle trajectory. **E.** By painting those *events* with increasing luminance in blue and those with decreasing luminance in black, the pattern of luminance change in each *cluster* can be visualised.

### 2.6 Twenty-two features used to characterise each individual particle

As the first attempt at aquatic particle observation, we decided on the following 22 characteristic features to characterise the shape, movement, and periodicity of each *cluster* (Fig. 3). Because the EVS cannot directly record the shape of a particle due to its non-frame-based nature, the shape must be calculated from the characteristics of the movement of a *cluster* (4 characteristics out of 22). Characteristics of the *cluster* movement are also described based on variances and differences of the *events* in a *cluster* (8 characteristics out of 22). To utilize EVS’s advantage of high-speed recording, millisecond-scale periodicity in *cluster* behaviours is examined (10 characteristics out of 22). Criteria for the calculation can be tuned to obtain appropriate information from the EVS output according to the research question.

**Figure 3.**
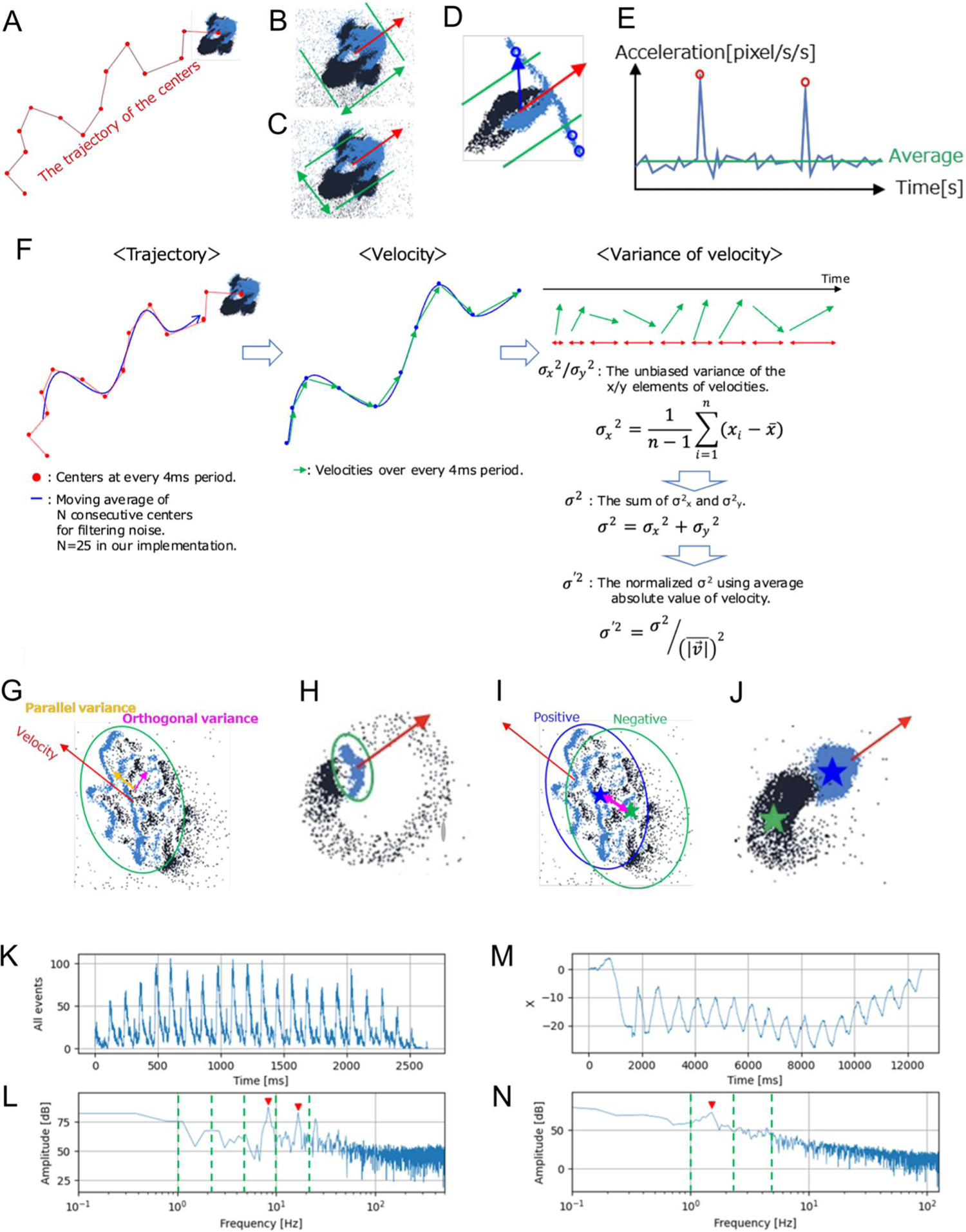
Twenty-two features calculated to characterise *clusters*. **A.** F01 Average velocity: The coordinates of all *events* in a *cluster* were averaged and defined as the centre of the *cluster*. The trajectories of the centres were segmented with a fixed period of time and the velocity of the *cluster* was calculated at each segment. The velocity values were averaged across all trajectories. **B, C.** F02 Body length, F03 Body width: Length and width of *cluster* were defined as the average number of pixels parallel or orthogonal to the velocity (F01), respectively. **D.** F04 Antenna indicator: The inner product of the velocity (F01) and the vector from the centre to an *event* extending out of the body width was accumulated across all trajectories and divided by its length (F02). **E.** F05 The number of large acceleration peaks: Acceleration was defined as the absolute value of the derivative of velocity (F01). Large acceleration peaks with values over five times greater than the average acceleration across all trajectories were counted. **F.** F06 Variance of velocity: For *clusters* with the average velocity over 0.025 [pxl/msec], those with the variance of velocity higher than 0.14 were classified as *Active* or *Dynamic particles*, while the rest were classified as *Beating* or *Passive particles*. In combination with F13-F22, these *clusters* can be classified into four groups: *Active, Dynamic, Beating* and *Passive particles*. **G, H.** F07–10 Parallel/Orthogonal variance of positive/negative *events:* The average variance parallel/orthogonal to the velocity of positive *event* coordinates (blue pixels in G, H) were calculated separately and each value was divided by its length/width resulting in F07/F08, respectively. The same calculations were used on negative *events* resulting in F09/F10 (black pixels in G, H). **I, J.** F11, 12 Parallel/Orthogonal distance ratio between positive and negative centres: The average distance between centres of positive and negative *events* was calculated and then divided by its length/width resulting in F11/F12, respectively. **K, L.** F13–18 FFT peaks of *event* increase: Temporal patterns of the increase of the positive, negative, and total *events* (positive and negative) of each *cluster* were analysed by Fast Fourier Transformation (FFT) with the time resolution of 1 msec. Rectangular window with the length of entire track duration was used as the time window function. For better classification of living organisms based on specific periodicity of particle behaviours, the entire frequency range of the power spectrum generated by FFT was divided into six segments; <1, <2.2, <4.6, <10, <22, and ≥22 Hz respectively, assigned from F13 to F18, respectively. Peaks with values greater than the background level by 15 dB in the power spectrum were counted. The peak indicates the presence of periodic movement of the *clusters* themselves. At each segment, the numbers of detected peaks were summed across all the FFT results of the positive, negative, and total *events*. **K.** An example of the increase of both the positive and negative *events* at every 1 msec of an *Artemia* nauplius larva showing a saw-toothed pattern. **L.** FFT analysis of the data shown in panel **K** detected two peaks (red arrowheads) that above the threshold. **M, N.** F19–22 FFT peaks of centre coordinates: Temporal patterns of the x/y coordinates of centres of the positive, negative, and total of positive and negative *events* of each *cluster* were analysed by FFT with the time resolution of 4 msec. A linear function was subtracted from the x/y coordinates so that the resultant x/y coordinates start and end at the coordinate of 0. Then, rectangular window with the length of entire track duration was used as the time window function. The entire frequency range was divided into four segments; <1, <2.2, <4.6, and ≥4.6 Hz respectively, assigned from F19 to F22. Peaks with values greater than the background level by 15 dB in the power spectrum were counted. The peak indicates the presence of periodicity in the positions of the *clusters*. At each segment, the numbers of detected peaks were summed across all the FFT results of the positive, negative, and total *events*. **M.** An example of x coordinates of the total of positive and negative *events* in the veliger of the limpet *N. fuscoviridis*. **N.** The result of FFT analysis of M showing a peak (red arrowheads).

#### F01 Average velocity

The coordinates of all the *events* in a *cluster* were averaged, the resulting averaged coordinate was defined as the ‘centre of the *cluster*’. The trajectory of the centres was segmented with a fixed period of time and the velocity of the *cluster* was calculated at each segment. The speed was averaged across all the trajectories (Fig. 3A).

#### F02-03 Body length and width

Length (F02) and width (F03) of each *cluster* were defined as the average number of pixels parallel/orthogonal to the velocity (F01), respectively (Fig. 3B, C).

#### F04 Antenna indicator

The inner product of the velocity (F01) and the vector from the centre to an *event* extending some distance away from the body width was accumulated across all the trajectories and divided by its length (F02) (Fig. 3D),

#### F05 The number of large acceleration peaks

Acceleration was defined as the absolute value of the derivative of velocity (F01). Large acceleration peaks with values over five times larger than the average acceleration across all the trajectories were counted (Fig. 3E).

#### F06 Variance of velocity

For *clusters* with average velocity of higher than 0.025 [pxl/msec], those with the variance of velocity over 0.14 were labelled as *Active* or *Dynamic particles*, while the rest were labelled as *Beating* or *Passive particles*. In combination with F13-F22 (see below), particles could be classified into four groups: *Active*, *Dynamic*, *Beating*, and *Passive particles* (Fig. 3F, see below).

#### F07-10 Parallel/orthogonal variance of positive/negative events

The average variance of positive *event* coordinates parallel/orthogonal to the velocity (blue pixels in Fig. 3G, H) were calculated separately and each value was divided by its length/width resulting in F07/F08, respectively. Same calculations were operated on negative *events* resulting in F09/F10 (black pixels in Fig. 3G, H).

#### F11-12 Parallel/orthogonal distance ratio between positive and negative centres

The average distance between centres of positive and negative *events* was calculated and then divided by its length/width resulting in F11/F12, respectively (Fig3. I, J).

#### F13-18 Fast Fourier Transformation (FFT) peaks of event increase

Temporal patterns of the increase of the positive, negative, and the total *events* (i.e. both positive and negative) of each *cluster* were analysed by Fast Fourier Transformation (FFT) with the time resolution of 1 msec. A rectangular window with the length of entire track duration was used as the time window function. For a better classification of plankton based on specific periodicity of particle behaviours, the entire frequency range of the power spectrum generated by FFT was divided into six segments; <1, <2.2, <4.6, <10, <22, and ≥22 Hz, which were respectively assigned to F13 to F18. At each segment, the numbers of detected peaks were summed across all the FFT results of the positive, negative, and total *events*. The peaks indicate the presence of periodic movement of the *particles* themselves. An example of the FFT analysis of the all *events* of a metanauplius larva of the brine shrimp *Artemia* is shown in Fig. 3K and L.

#### F19-22 FFT peaks of centre coordinates

Temporal patterns of the x/y coordinates of centres of the positive, negative, and total *events* of each *cluster* were analysed by FFT with the time resolution of 4 msec. A linear function was subtracted from the x/y coordinates so that the resultant x/y coordinates start and end at the coordinate of 0. Then, a rectangular window with the length of the entire track duration was used as the time window function. The entire frequency range was divided into four segments; <1, <2.2, <4.6, and ≥4.6 Hz, assigned respectively to F19-F22. At each segment, the numbers of detected peaks were summed across all the FFT results of the positive, negative, and total *events*. The peaks indicate the presence of periodicity in the positions of the *particles*. An example of the FFT analysis of x coordinates of the all *events* of a veliger larva of the true limpet *Nipponacmea fuscoviridis* is shown in Fig. 3M and N.

For *clusters* with no peak detected in the FFT analyses (i.e., F13-F22 values were all 0), those with F6 values of 0.14 or smaller were placed in the *Passive particle* category and those with F6 values over 0.14 were categorised as *Dynamic particle*. For *clusters* where one or more peaks were detected in the FFT analyses (i.e., one or more of F13-F22 was ≥1), those with F6 values equal or less than 0.14 were placed in the *Beating particle* category, those with F6 values over 0.14 were considered to be in the *Active particle* category. The flow diagram of this classification of particles into the four categories is shown in Fig. 4.

**Figure 4.**
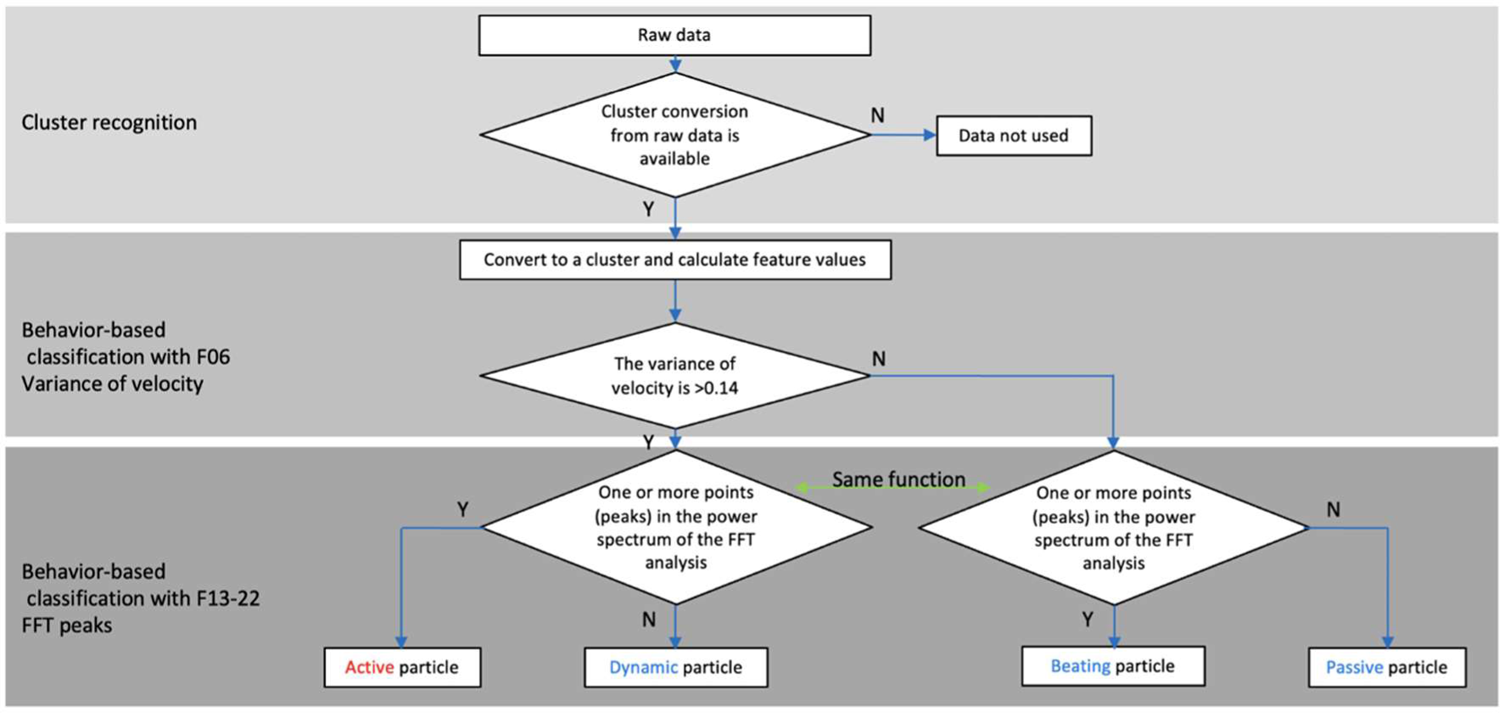
Particle classification flow. *Clusters* are mechanically classified from the raw data into the four categories of particles: *Active*, *Dynamic*, *Beating*, and *Passive particles* using this flow.

### 2.7 Machine learning for plankton classification

To extract feature values from each species, we first assigned IDs to each *cluster* identified through data analysis. Then, using the frame-based images recorded simultaneously with the EVS system, we classified each *cluster* into five categories: nauplius larva of *Parvocalanus crassirostris* (Pcra_N), copepodite and adult of P. *crassirostris* (Pcra_C), *N. fuscoviridis* (Nfus), *Patiria pectinifera* (Ppec), and passive particles (Passive) including non-swimming plankton, and dust. After annotation of *clusters* recorded by the EVS using the frame-based records, we selected representative individuals for each species from the entire data based on the clarity of the *cluster* shape and independence (i.e., considered to be independent when there is no collision with other particles in the 2D view).

With the EVS output of three species observation, the 22 features were calculated and fed into the fully-connected three-layer neural network (Wasserman and Schwartz, 1988) for the classification of each *cluster* into Pcra_N, Pcra_C, Nfus, Ppec, and Passive (Fig. 7). Nauplius larvae of *Paryocalanus*. *crassirostris* (Pcra_N) and copepodites of *Paryocalanus crassirostris* (Pcra_C) were distinguished for the ease of analysis and learning due to the significant difference in their movement and shapes to each other. 138 *cluster* were used as teaching data, while 119 *cluster* were used as test data for validation of the analysis.

**Figure 5.**
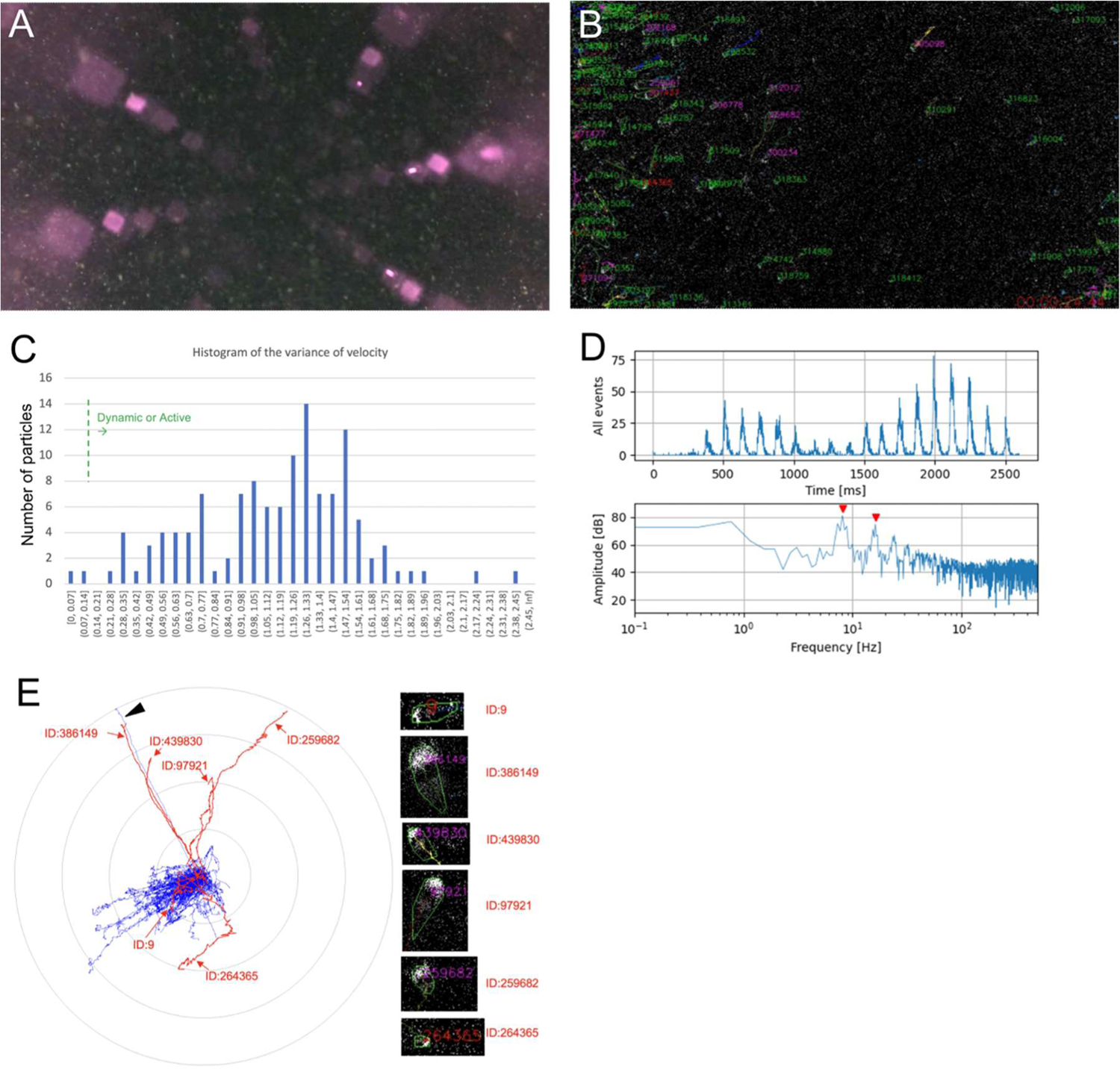
Results from Lake Biwa with white light. **A.** A frame from video recorded by a conventional frame-based video camera. Numerous particles were observed (whitish dots). The purple light is the light reflected on the surface of the glass sphere. **B.** A snapshot of the same time as **A** in the result of analysing EVS recordings. Many *clusters* were detected. **C.** A histogram of variance of velocity (F06) for a given 40 seconds. X-axis is variance of velocity and Y-axis is number of *clusters*. Most of the clusters exhibited values higher than the threshold for them to be classified as *Passive* or *Beating* (0.14, green dashed line). **D.** The result of the FFT analysis of *cluster* ID: 386149, which was classified into *Active particle*. Upper: increment of all *events*, lower: power spectrum. The power spectrum showing two peaks at 8.1 Hz and 16 Hz (red arrowheads). **E.** A radar chart of the *cluster* trajectories detected during the 40 seconds. The six clusters classified into *Active* are shown in red and the other in blue. The *Active particles* were moving in various directions, while most of the other clusters were moving to the lower left. A *cluster* (arrowhead) was classified into *Passive*, despite the almost same trajectory with an *Active particle*, ID: 386149.

**Figure 6.**
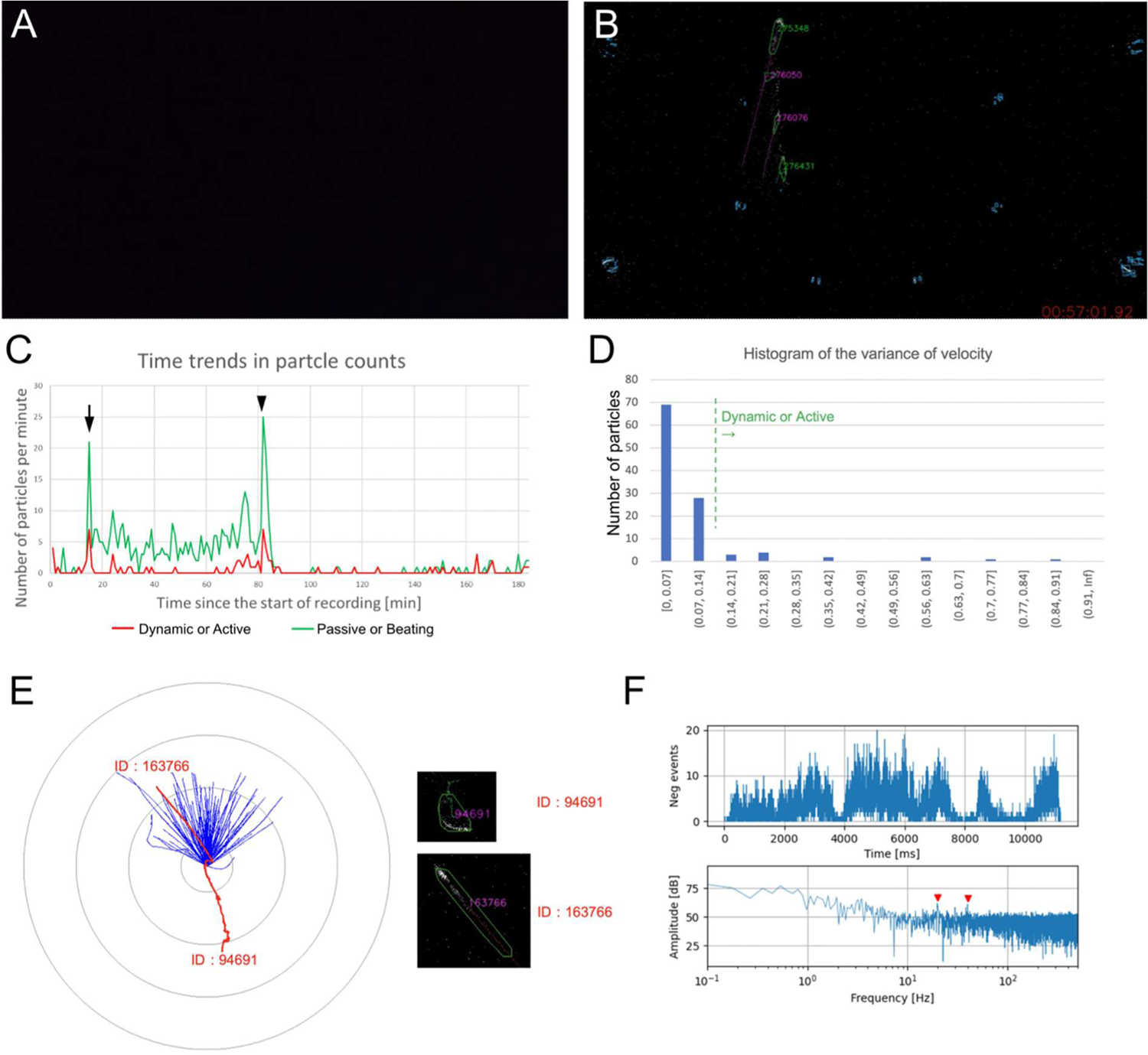
Results from the deep sea with infrared light. **A.** A captured image from a movie recorded by the frame-based video camera, which was unable to capture any image due to low light. **B.** A snapshot of the same time as A from the EVS showing the result of analysing its recordings. Even under very faint infrared-light condition, the EVS was able to detect *events* and identify *clusters*. **C.** Time-series changes in the number of *Dynamic* or *Active* (red) and *Passive* or *Beating* (green) *particles* during the survey by ROV. Many *clusters* were detected at the time of the ROV arriving on sea bottom (arrow) and started to move (arrowhead). **D.** A histogram of variance of velocity (F06) for a given 20 min at the seafloor. X-axis is variance of velocity and Y-axis is number of *clusters*. Almost all *clusters* were classified as *Passive* or *Beating* (threshold 0.14, green dashed line). **E.** A radar chart of the *cluster* trajectories detected during the same 20 min. The two *clusters* classified into *Active particle* are shown in red and the others in blue. **F.** The result of the FFT analysis of *cluster* ID: 94691, which was classified into *Active particle*. Upper: increment of negative *events*, lower: power spectrum. The power spectrum showing two peaks at 20 Hz and 40 Hz (red arrowheads).

**Figure 7.**
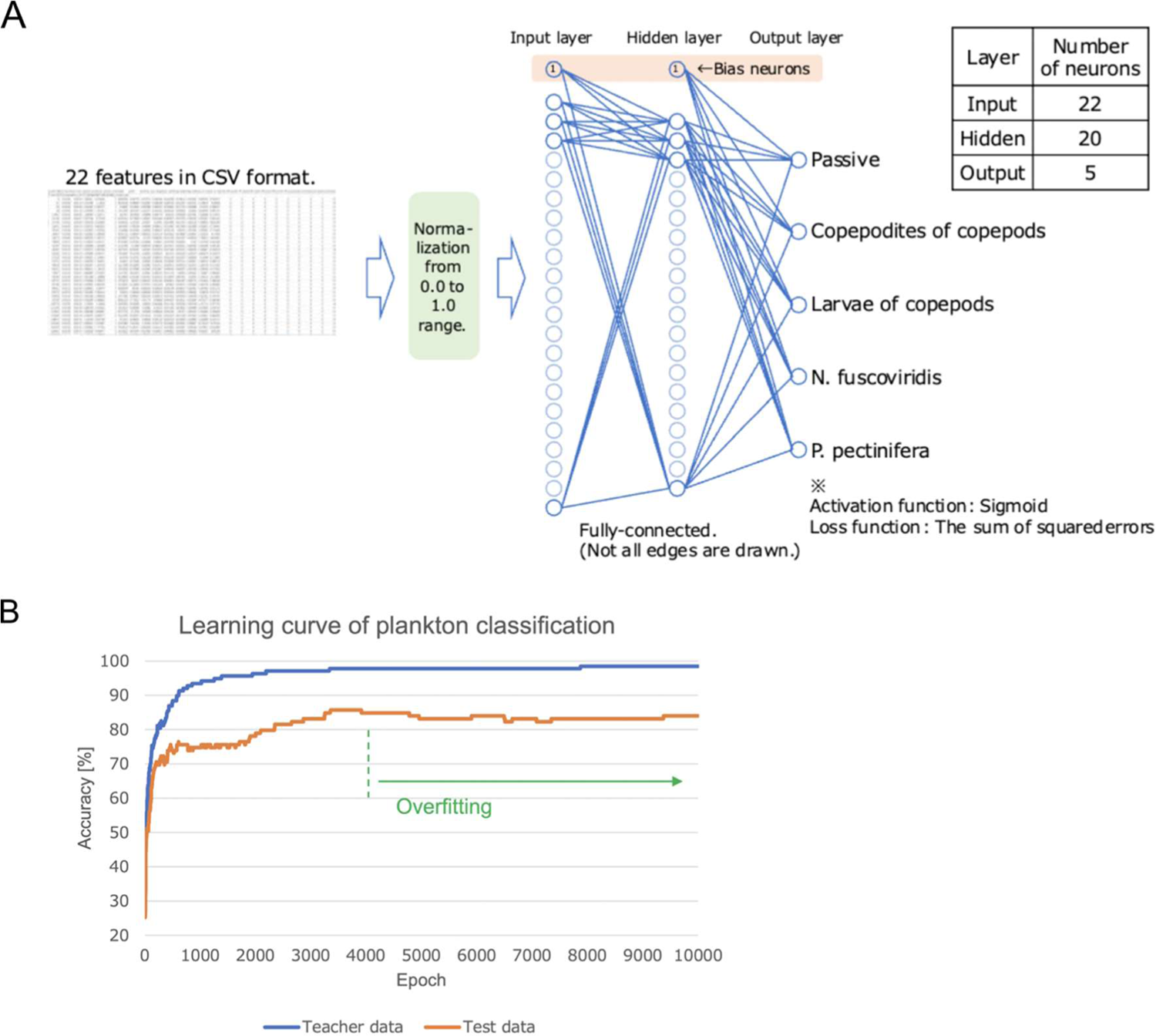
Machine learning process used for plankton classification. **A.** The 22 pre-determined characteristic features were fed into a fully-connected tri-layer neural network for the classification of each *cluster* into nauplius larva of *Parvocalanus crassirostris* (Pcra_N), copepodite of *Parvocalanus crassirostris* (Pcra_C), veliger larva of *N. fuscoviridis* (Nfus), bipinnaria larva of *Patiria pectinifera* (Ppec), and passive particle. **B.** Learning curve of the plankton classification. The x-axis shows the number of iterations in which all the teacher data were trained (epoch), and the y-axis shows the percentage of correct responses when the teacher data (blue) and the test data (orange) were judged at that number of epochs. The correct response rate for the teacher data increased with the number of training sessions. When the test data was judged, the correct response rate increased at almost the same rate as the teacher data up to about 300 epochs. After that, the correct response rate of test data was lower than that of the teacher data, but the rate gradually increased to a maximum of 86%. On the other hand, after about 4000 epochs, the correct response rate either plateaued or declined, due to overfitting to the teacher data.

## 3 Results and Discussion

### 3.1 Evaluation 1: Detection of active particles in passive particle storms examined through *in situ* observation

#### Lake Biwa

Raw EVS outputs during the Lake Biwa observation were analysed to quantitatively evaluate the behaviours of particles in the natural lake water. During the observation period when the conventional frame-based camera captured intensive movements of numerous particles (Fig. 5A), the EVS system also detected numerous *clusters* (Fig. 5B, Supplementary Movie S1). To give an example, the 125 *clusters* identified during a 40-sec recording at 11:44 JST were categorised based on the particle classification flow (Fig. 4). Among the 125 *clusters*, 123 *clusters* had variances of velocity (F06) greater than the threshold of 0.14, and only two *clusters* had values less than the threshold (Fig. 5C). According to the next step of the particle analysis flow (Fig. 4, bottom), those 123 *clusters* with F06 over 0.14 were classified into 6 *Active* and 117 *Dynamic*, and the other two *clusters* were classified into 1 *Beating* and 1 *Passive*. The power spectrum for an *Active particle* (ID: 386149) revealed that the particle exhibited periodic motions at the frequencies of 8.1 Hz and 16 Hz (Fig. 5D). Such integer multiple frequency peaks are considered to be harmonics. Periodicities were also found in four of the other five *Active particles*, which were between a narrow range of 5.8 Hz and 7.7 Hz (power spectrum not shown), while the last *Active particle* (ID: 9) had a periodicity of 1.3 Hz. The periodicity in *cluster* motion suggests that the specific periodicities correspond to characteristics of swimming behaviour of species.

Furthermore, the predominance of *Dynamic particles* compared to other *particle* types are unlikely to represent a collectively-synchronized behaviour of the whole plankton community, but instead likely results from the strong water flow and the shaking of suspended particles, as well as the camera system itself shaking due to the instability of the rope-anchored COEDO. This means that the F06 feature (variance of velocity) calculated from the dataset acquired by a floating camera system in a system impacted by water current is likely not an appropriate characteristic for distinguishing living particles from non-life. Alternatively, improving the mathematical processing to eliminate components exhibiting synchronized motion, found in the great majority of *clusters*, will help to correct for the effects of camera motion in *in situ* observations. This improvement on the data processing was not included in the analysis software used in the present study, but will be considered in the future.

Results based on the automated classification analyses were verified by visual observation of pseudo-frame-based images, simulated using a 40-sec-integrated subset of the EVS data. While almost all the *clusters* moved in overall the same direction, some *clusters* (e.g., ID: 386149) appeared to be actively moving against the direction of other *clusters* (Fig. 5E, Supplementary Movie S1). Cumulative trajectories of each *cluster* revealed distinct moving behaviours of the *Active particles*, as classified by our software (Fig. 5E). The cumulative trajectories, however, revealed that the present automated classification sometimes misclassified *clusters* which appeared to be swimming organisms into non-*Active particles*. For example, a *cluster* swimming in the same direction as an *Active particle*, ID: 386149, was classified into *Dynamic particle* because of the lack of frequencies (Fig. 5E, arrowhead). The pseudo-frame-based video image also indicated that the distinct periodicity of the *Active particle* (ID: 9) was a calculation artifact, due to frequent collisions/separations with neighbouring *clusters*. These results suggest that further mathematical optimization of *cluster* recognition, characteristic features developed to classify *clusters*, and criteria for *cluster* classification is required to improve the accuracy of automatic identification of particle types in aquatic environments.

#### Deep-sea Suruga Bay

During the descent to the seafloor in the deep-sea observation at Suruga Bay, the EVS captured only a few *clusters* in the deep-sea water column. At the time of the ROV arriving on bottom at 10:20 JST (Fig. 6C, arrow), the number of *clusters* identified by the EVS increased, suggesting the resuspension of sedimentary particles and/or escape behaviour of benthic animals due to seafloor disturbance by the ROV. These *clusters* were classified into not only *Passive* or *Beating*, but also *Dynamic* or *Active* ones, based on the criteria mentioned above. Through the entire 180-minutes of EVS recording, the number of *Dynamic* or *Active particles* were approximately 10 times fewer than *Passive* or *Beating clusters* (Fig. 6D). Cumulative trajectories of each *cluster* as well as the pseudo-frame-based video, reconstructed from a 20-min subset of the EVS recording (from 10:14 to 10:34), showed that almost all *clusters* moved upwards (Fig. 6E, Supplementary Movie S2). Based on the particle classification flow (Fig. 4), two *clusters* were classified into *Active particles* during those 20 minutes. One of the two (ID=94691) showed a distinct trajectory from the others, suggesting swimming behaviour against the flow (Fig. 6E). The power spectrum of the negative *events* of this *Active particle* (ID=94691) exhibited periodic motions at the frequencies of 20 Hz and 40 Hz, while probably being harmonics (Fig. 6F). The conventional frame-based camera of the EVS system (GoPro Hero 8) was unable to capture clear visuals from 100 m below the sea surface, due to insufficient sensitivity of sensor elements (Fig. 6A). In contrast, EVS could detect particles under the very faint infrared LED light (Fig. 6B).

### 3.2 Evaluation 2: Distinction among different species based on particle behaviours examined through *in vitro* observation of cultivated plankton

#### Machine learning for plankton classification

The correct response rate for the teaching dataset increased with the number of training sessions (Fig. 7B, blue). When the test dataset was run in the trained neural network, the correct response rate increased at almost the same rate as the teaching dataset up to about 300 epochs (Fig. 7B, orange). After that, the correct response rate of the test dataset was lower than that of the teaching dataset, but the rate gradually increased to a maximum of 86%. On the other hand, after about 4000 epochs, the correct response rate either plateaued or declined, due to overfitting (Lawrence, Giles and Tsoi, 1997) to the teaching dataset. The high accuracy achieved by a relatively small number of teaching dataset (a total of 138) means that the EVS data contained enough amount of information from which meaningful features of each species’ movement and shape could be calculated. It also means that the 22 characteristic features used in this study are meaningful and sufficient for classifying a variety of living organisms and *Passive particles*. Note that our mathematical analysis of the EVS dataset recognizes living organisms without any motion as *Passive particles*. The rapid movement of *clusters* and intersections between *cluster* trajectories in a 2D space often interfere with the tracking accuracy of each *cluster* when *clusters* were automatically identified by the software. The ultra-high fps of the EVS system, however, means detailed information is available for each *cluster* until just before the intersection, allowing for the continuous tracing of *cluster* by manual curation and therefore considerably decreasing the interference even when a large number of particles are present.

#### Species-specific FFT peaks

We performed two types of frequency analyses: “event FFT peak” (F13-18) and “coordinate FFT peak” (F19-F22). In the limpet *Nipponacmea fuscoviridis*, 33.3% and 23.3% of individuals exhibited “event FFT peak” and “coordinate FFT peak”, respectively (Fig. 8A, Nfus). Nauplius larvae of the copepod *Parvocalanus crassirostris* showed low percentages of both “event FFT peak” and “coordinate FFT peak”, at 6.8% (Fig. 8A, Pcra_N). Copepodites and adults of *Parvocalanus crassirostris* also showed low percentages of FFT peak, 0% in “event FFT peak” and 6.5% in “coordinate FFT peak” (Fig. 8A, Pra_C). The result showing few *clusters* in this species exhibited frequencies in the EVS signals may reflect that this species does not exhibit much periodic behaviour. In the sea star *Patiria pectinifera*, while 45.5% of individuals exhibited “event FFT peaks”, no individual exhibited “coordinate FFT peaks” (Fig. 8A, Ppec). Passive particles displayed a low percentage (5.9%) of both “event FFT peaks” and “coordinate FFT peaks” (Fig. 8A, Passive). Metanauplius larvae of the brine shrimp *Artemia* had high percentages of individuals exhibiting both FFT peak types, 90.2% with “event FFT peaks” and 95.1% with “coordinate FFT peaks” (Fig. 8A, Artemia). We also analysed the FFT peaks of the two phytoplankton species studied. *Chattonella marina* swims with two flagella, and in this species 21.7% of individuals observed exhibited “event FFT peaks” and 4.3% had “coordinate FFT peaks” (Fig. 8A, Cmar). *Heterosigma akashiwo* also swims with two flagella, and 13.3% of individuals exhibited both “event FFT peaks” and “coordinate FFT peaks” (Fig. 8A, Haka). Taken together, these results indicate that plankton swimming in the water column do not always exhibit either FFT peaks, but the presence/absence of the FFT peaks provides a strong indicator for classifying actively swimming and passive particles.

**Figure 8.**
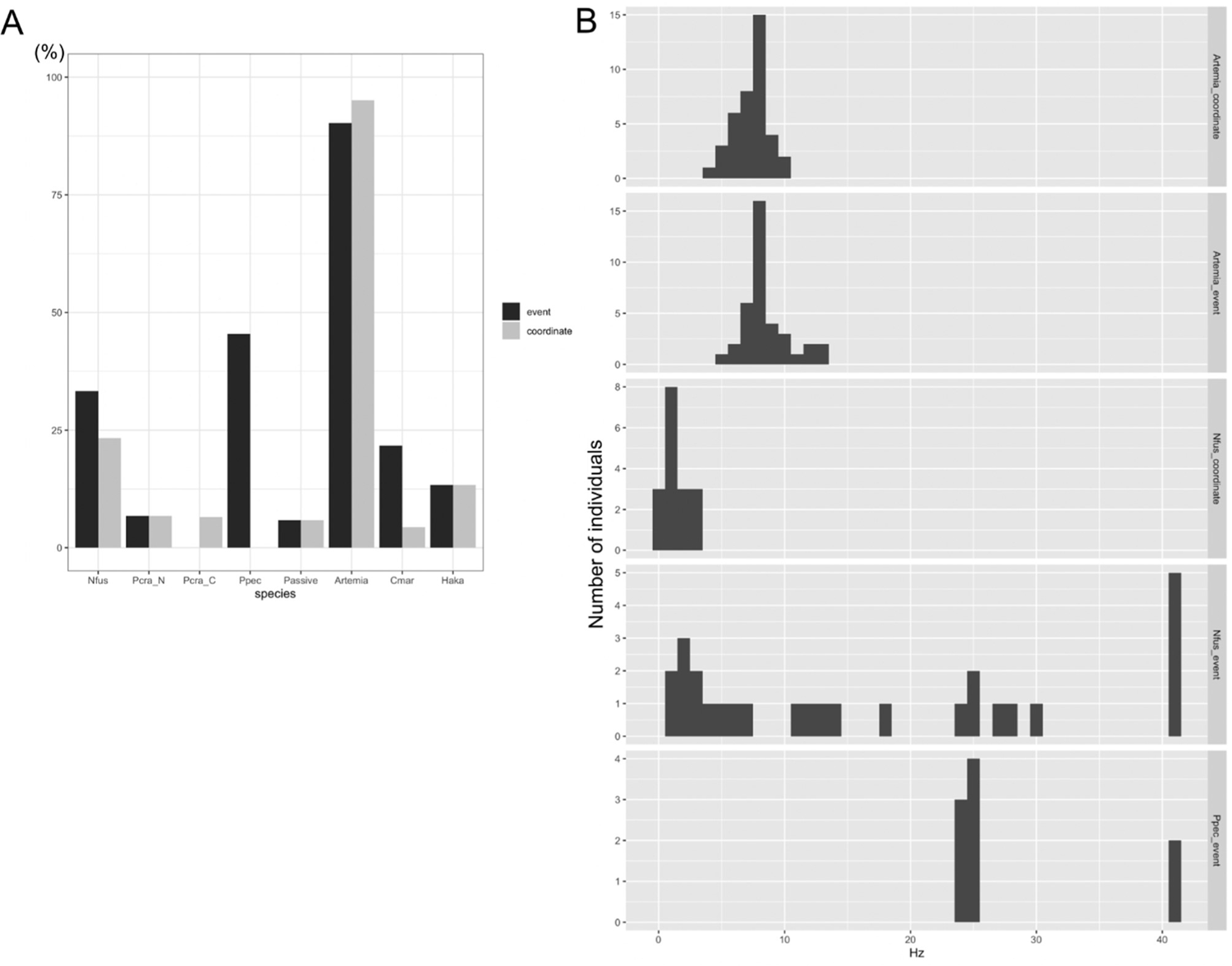
Species specificity for FFT peaks. **A.** A graph showing the percentages of individuals with FFT peaks at the increase in the number of *events* or the x/y coordinates of centres of the *cluster*. **B.** Histograms of FFT peak values for each species showing that each species has a characteristic frequency.

We found that some plankton species tended to possess characteristic frequencies. The FFT analysis of metanauplius larvae of the brine shrimp *Artemia* showed that they exhibited “event FFT peaks” at around 7 Hz among individuals (Fig. 8B, Artemia_event). The increase and decrease of *events* in this 7 Hz cycle was consistent with the anteroposterior movement of the appendages in the swimming behaviour of *Artemia* (Supplementary Movie S3). The 7 Hz cycle of the appendage movement resulted in the “coordinate FFT peak” being distributed around 7 Hz in this stage of larvae (Fig. 8B, Artemia_coordinate). Veliger larvae of the limpet *N. fuscoviridis* tended to show “coordinate FFT peaks” at about 1.5 Hz (Fig. 8B, Nfus_coordinate). This periodicity of the central coordinates reflects the spiral swimming pattern of these veliger larvae. On the other hand, there was no clear trend in the distribution of “event FFT peak” in the veliger larvae of *N. fuscoviridis* (Fig. 8B, Nfus_event). Most larvae of the sea star *Patiria pectinifera* exhibited “event FFT peaks” at approximately 24.6 Hz (Fig. 8B, Ppec_event); though it is unclear what behaviour this links to, it may be related to ciliary beating in the ciliary band. These results indicate that the EVS system and subsequent analyses using our software can unveil characteristics of rhythmic behaviours of organisms. The extremely high temporal resolution of the EVS enabled FFT analysis at a high frequency range, leading to the detection of “event FFT peaks” at over 40 Hz with *N. fuscoviridis* and *Patiria pectinifera* (Fig. 8B). This cannot be achieved with conventional cameras with only typically 50/60Hz temporal resolution, whose frequency range would be limited to about 25/30Hz for FFT analysis due to the sampling theorem.

## 4. Concluding remarks

Our tests of the EVS system *in situ* in a lake and a deep-sea environment as well as *in vitro* in controlled laboratory aquaria showed that the system is capable of identifying and classifying the movement of particles in aquatic environments. Ultra-high-speed movement tracking and wide dynamic range of the EVS system allow us to quantify the high-speed millisecond-order rhythmic motions of plankton under weak infrared LED lighting, as well as the dynamics of plankton swimming among numerous passive particles in relatively turbid lake water. The 22 characteristic features developed for the classification and analysis of the EVS output are sufficiently functional to distinguish particles based on their behaviour, as seen by *in situ* observations and confirmed by laboratory experiments. The machine learning of particle classification achieved at most 86% accuracy.

These results collectively suggest a great potential for the EVS in carrying out *in situ* observations of plankton and/or particles in aquatic ecosystems to understand their biology and to monitor their responses to climate change. The system is sufficiently compact to integrate with buoys or floats like ARGO (Roemmich et al., 2019) or autonomous underwater vehicles for the sensing of deep-sea plankton (Ohman et al., 2019), followed by edge computing (i.e. data processing directly on the device) for analyses and classification. The EVS workflow performed in the present study can be applied to analyse the behaviour of microscopic organisms with a high temporal resolution, as well as cellular-level motion analysis, such as sperm motility.

Furthermore, the EVS has wider applications to other larger organism groups. For example, breakthroughs in tracking devices and satellite communication now allow for the so-called bio-logging approach to incorporate video data (Watanabe & Takahashi, 2013), and there are year-long video monitorings of underwater communities (Aguzzi et al., 2019). The EVS can be added to these systems or replace conventional video cameras in order to gain more in-depth and automated understanding of organismal behaviour. In addition, many larger animals outside of aquatic environments also exhibit rapid motions, such as the vocal cord during sound production. Recently, biomechanical studies of vocal fold vibration in laryngeal sound production and pathophysiological evaluation of voice disorders have been carried out using high-speed frame-based video analyses (Schützenberger et al., 2016), but the EVS offers a more superior method for these applications due to its ultra-high fps.

Observing natural phenomena using the EVS is a newly emerging method and will benefit greatly from future improvements, especially to the data analyses. For example, more appropriate and specific characteristic features to represent the particle behaviours and classification criteria could be developed. Three-dimensional behaviour of the target object can also be reconstructed by combining multiple EVS systems. The accuracy of particle classification through machine learning can be improved by further iterations with more diverse teaching data and more suitable feature settings. These improvements will be the foci of future studies.

## Supporting information

Supplementary_Information_biorxiv

## Acknowledgement

The COEDO operation at Lake Biwa was supported by the Japanese Council for Science, Technology, and Innovation (CSTI), Cross-ministerial Strategic Innovation Promotion Program (SIP) “Innovative Technology for Exploration of Deep-Sea Resources” (Lead agency: JAMSTEC). We thank the captain, crew, and on-board scientists of R/V *Kairei* during cruise KR21-17C, and the operation team of ROV *Kaiko*. We also thank the staff of marine biological stations of Research Center for Marine Biology of Tohoku University and Marine and Coastal Research Center of Ochanomizu University for collecting adult sea stars used in this study. This work was supported by MEXT’s “Advancement of Technologies for Utilizing Big Data of Marine Life” Grant Number JPMXD1521473834.

## Author contribution

S.T., N.M. and S.K. conceived and designed the project. N.M., Y.M., Y.K. and A.Y. collected and maintained organisms. S.T. and H.S. developed hardware. H.S. developed software. S.T. and N.M. performed laboratory experiments. N.M. performed cluster annotation for machine learning. S.T., H.S. and S.K. performed *in situ* observations. N.M and C.C. performed biological interpretation. S.T., N.M., H.S., C.C. and S.K. interpreted the data and draft the paper. All authors contributed to the final paper.

