## Supplementary_Information_biorxiv for "Millisecond-scale behaviours of plankton quantified *in situ* and *in vitro* using the Event-based Vision Sensor (EVS)"

### Milli-second-scale behaviors of aquatic life quantified by intelligent event-based vision sensor

#### Legends of supplementary movies

**Supplementary Movie S1.**

**Supplementary Movie S2.**

**Supplementary Movie S3.**

#### Supplementary Figures

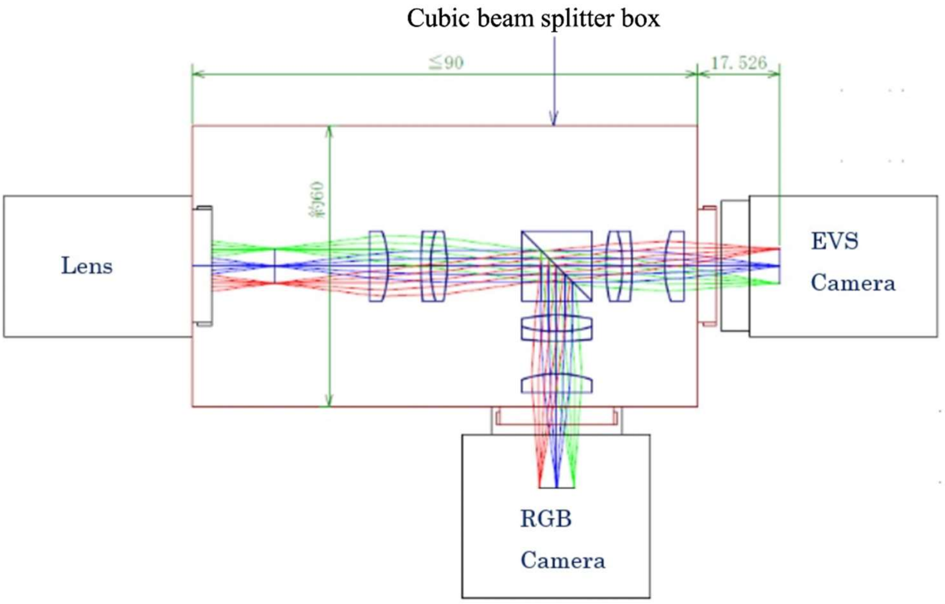


**Supplementary Figure S1.** The optical axis diagram of the cubic beam splitter box (ELIOTEC CORP.)

A B

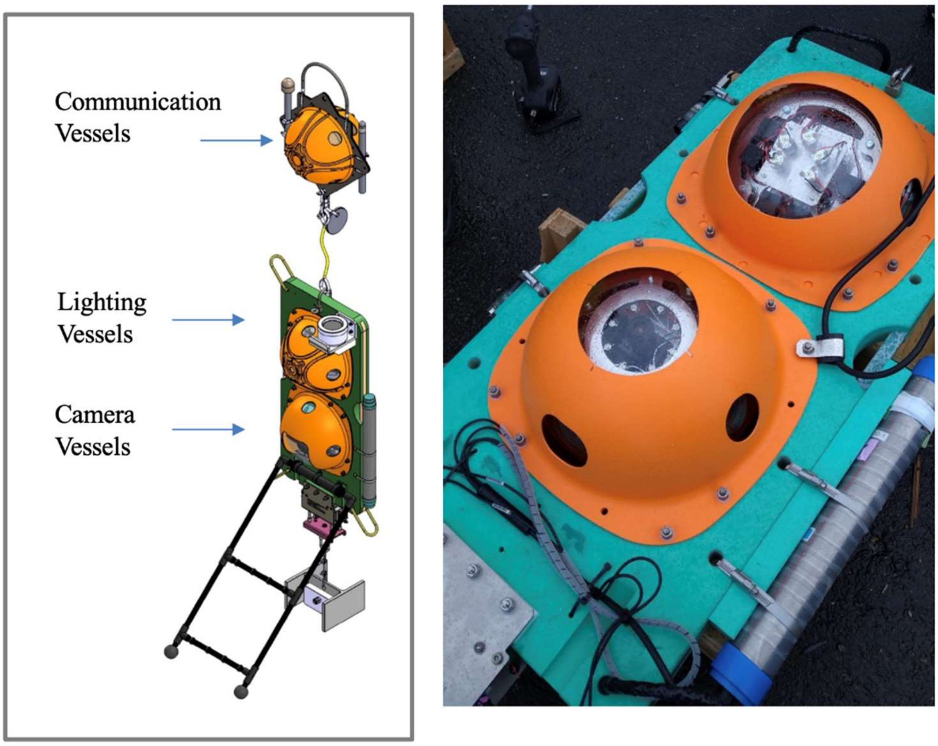


**Supplementary Figure S2. COEDO in-situ observation system. A.** A schematic image of COEDO system. **B.** A photograph of the light and camera spheres.

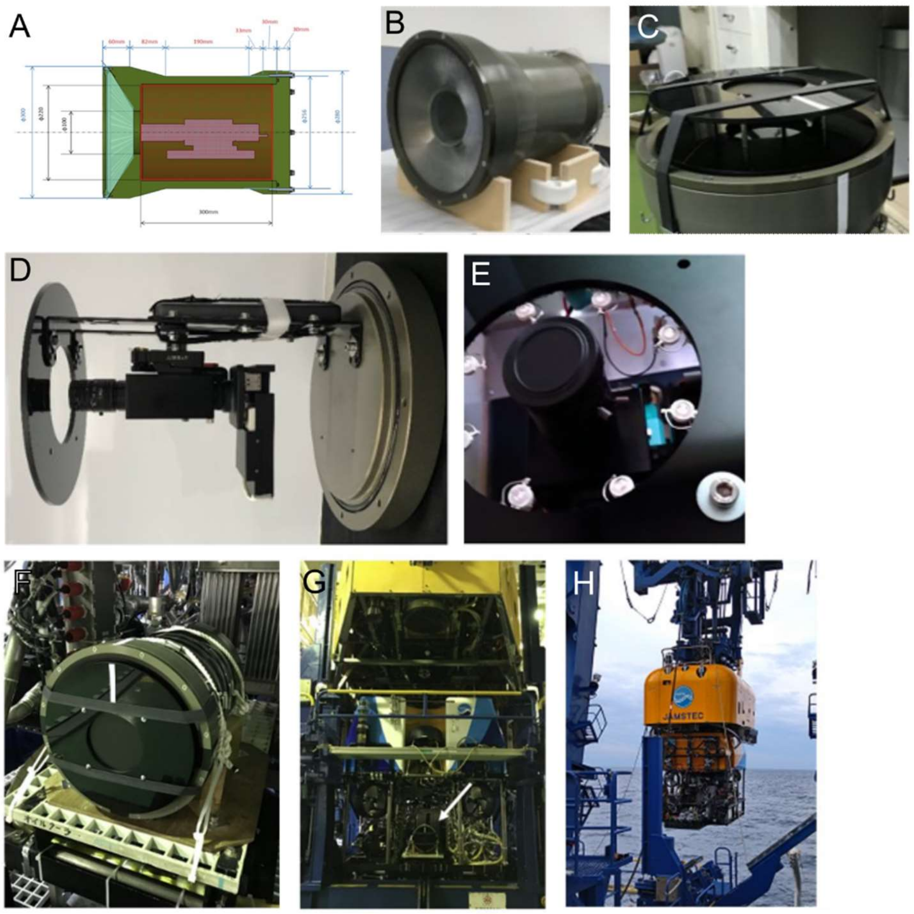


**Supplementary Figure S3. 3500m-class underwater camera housing for in situ observation. A.** A schematic diagram of the camera housing showing that the window of the housing has a 100 mm diameter, shieded by a 50 mm thick transparent acrylic plate (in the shape of a cone). **B.** A photograph of the camera housing. **C.** A close-up view of the window showing a shielding wall limiting the depth of the field of view. **D.** A photograph showing how the EVS prototype camera is fixed in the housing. **E.** A photograph showing 850 nm LED illumination postioned around the camera lens. **F–H.** Photographs show that the housing containing the EVS is fixed to the rear of the ROV Kaiko.

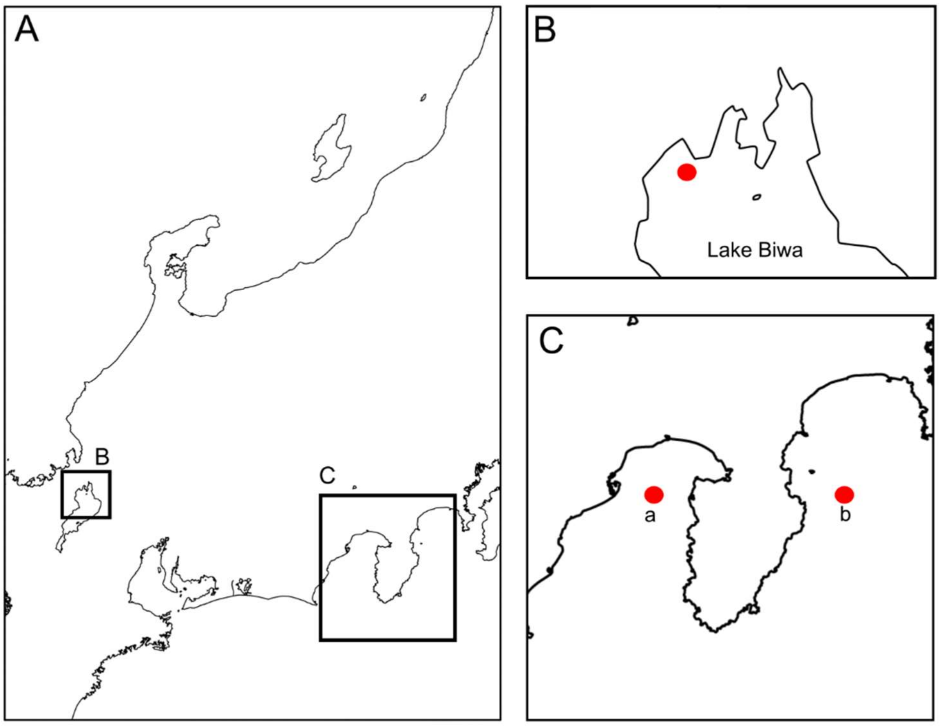


**Supplementary Figure S4. Maps showing the reserach sites of the present study. A.** A map of middle part of Japan showing the two research sites B (Lake Biwa) and C (Suruga Bay and Sagami Bay). **B.** A red dot showing the site where the COEDO system deployed. **C.** Red dots a (Suruga Bay) and b (Sagami Bay) showing the research sites using the ROV Kaiko Mk-IV.

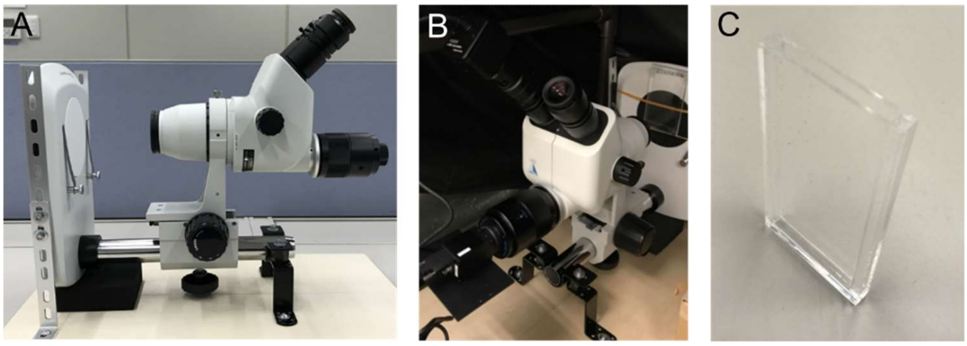


**Supplementary Figure S5. Imaging method in a laboratory condition. A.** A stereo microscope tilted 90 degrees and fixed. **B.** An EVS camera and an RGB camera (HOZAN USB Camera L-835) are connected to the microscope. **C.** A thin chamber for plankton observation.

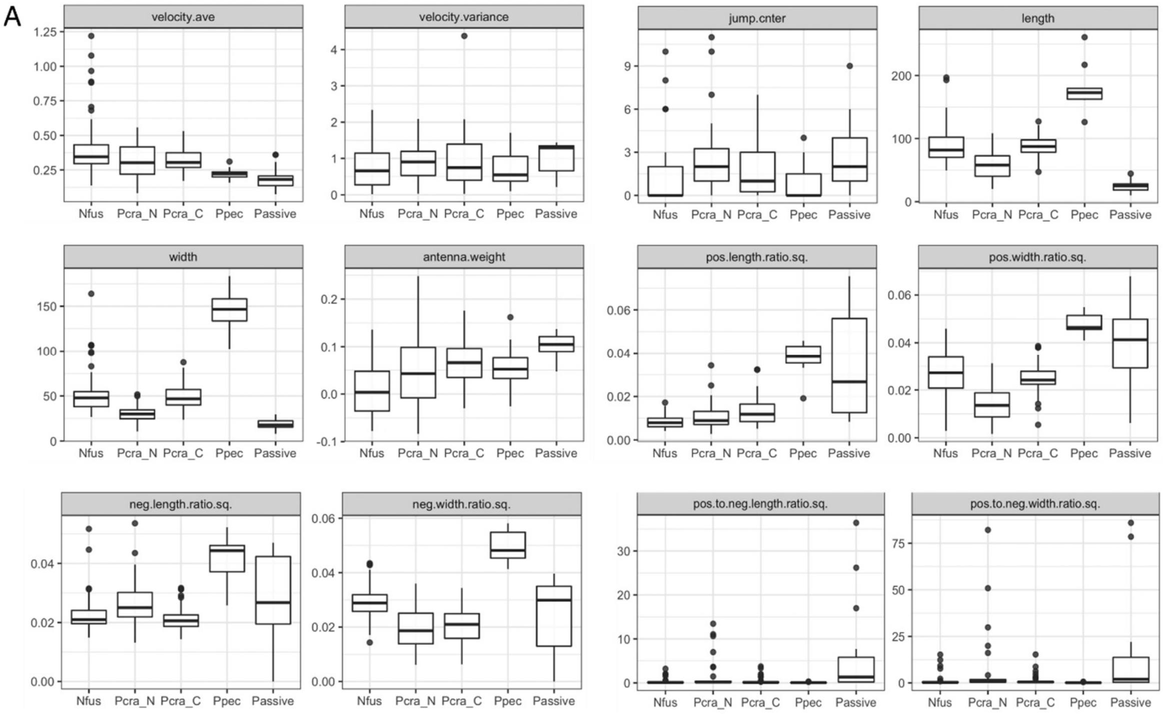


**Supplementary Figure S6.**

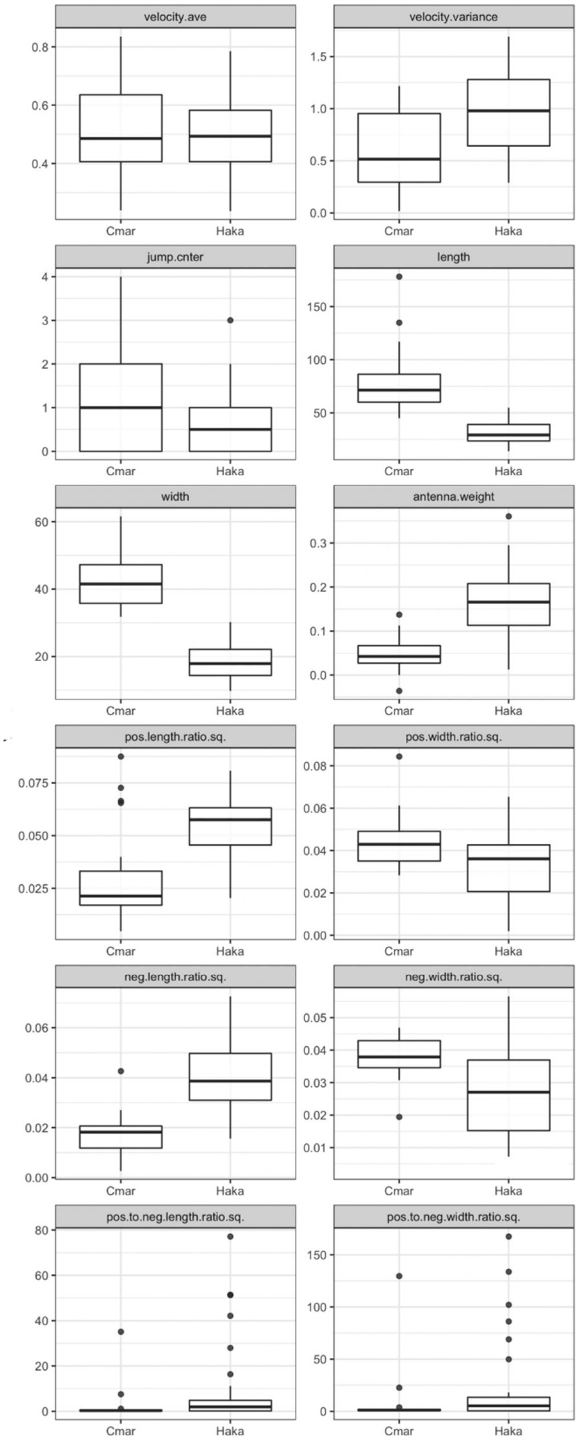


**Supplementary Figure S7.**
